# Efficacy of Parainfluenza Virus 5 (PIV5)-vectored Intranasal COVID-19 Vaccine as a Single Dose Vaccine and as a Booster against SARS-CoV-2 Variants

**DOI:** 10.1101/2022.06.07.495215

**Authors:** Ashley C. Beavis, Zhuo Li, Kelsey Briggs, María Cristina Huertas-Díaz, Elizabeth R. Wrobel, Maria Najera, Dong An, Nichole Orr-Burks, Jackelyn Murray, Preetish Patil, Jiachen Huang, Jarrod Mousa, Linhui Hao, Tien-Ying Hsiang, Michael Gale, Stephen B. Harvey, S. Mark Tompkins, Robert Jeffrey Hogan, Eric R. Lafontaine, Hong Jin, Biao He

## Abstract

Immunization with severe acute respiratory syndrome coronavirus 2 (SARS-CoV-2) vaccines has greatly reduced coronavirus disease 2019 (COVID-19)-related deaths and hospitalizations, but waning immunity and the emergence of variants capable of immune escape indicate the need for novel SARS-CoV-2 vaccines. An intranasal parainfluenza virus 5 (PIV5)-vectored COVID-19 vaccine CVXGA1 has been proven efficacious in animal models and blocks contact transmission of SARS-CoV-2 in ferrets. CVXGA1 vaccine is currently in human clinical trials in the United States. This work investigates the immunogenicity and efficacy of CVXGA1 and other PIV5-vectored vaccines expressing additional antigen SARS-CoV-2 nucleoprotein (N) or SARS-CoV-2 variant spike (S) proteins of beta, delta, gamma, and omicron variants against homologous and heterologous challenges in hamsters. A single intranasal dose of CVXGA1 induces neutralizing antibodies against SARS-CoV-2 WA1 (ancestral), delta variant, and omicron variant and protects against both homologous and heterologous virus challenges. Compared to mRNA COVID-19 vaccine, neutralizing antibody titers induced by CVXGA1 were well-maintained over time. When administered as a boost following two doses of a mRNA COVID-19 vaccine, PIV5-vectored vaccines expressing the S protein from WA1 (CVXGA1), delta, or omicron variants generate higher levels of cross-reactive neutralizing antibodies compared to three doses of a mRNA vaccine. In addition to the S protein, the N protein provides added protection as assessed by the highest body weight gain post-challenge infection. Our data indicates that PIV5-vectored COVID-19 vaccines, such as CVXGA1, can serve as booster vaccines against emerging variants.

**Importance:** With emerging new variants of concern (VOC), SARS-CoV 2 continues to be a major threat to human health. Approved COVID-19 vaccines have been less effective against these emerging VOCs. This work demonstrates the protective efficacy, and strong boosting effect, of a new intranasal viral-vectored vaccine against SARS-CoV-2 variants in hamsters.

## Introduction

Severe acute respiratory syndrome coronavirus 2 (SARS-CoV-2) first emerged in Wuhan, China in December 2019 [1]. Since then, it has spread globally, infected more than 519 million people, and caused at least 6 million deaths (https://covid19.who.int). SARS-CoV-2 initially infects the upper respiratory tract epithelium [2] but can progress to the lower respiratory tract and cause pneumonia and acute respiratory distress syndrome (ARDS) [3]. Since the beginning of the 2019 coronavirus disease (COVID-19) pandemic, numerous SARS-CoV-2 variants have emerged. The World Health Organization (WHO) defines a SARS-CoV-2 variant of concern (VOC) as a variant that affects virus transmissibility and COVID-19 epidemiology, increases virulence and pathogenicity, or decreases the effectiveness of COVID-19 vaccines (immune escape). Current VOCs include delta and omicron, while previously circulating VOCs include alpha, beta, and gamma (https://www.who.int/activities/tracking-SARS-CoV-2-variants).

The global spread of SARS-CoV-2 prompted rapid development of prophylactic vaccines. Currently, three vaccines are approved for use in the United States. The vaccines developed by Pfizer and Moderna are based on mRNA technology, while the vaccine produced by Johnson & Johnson (J&J) utilizes a human adenovirus type 26 vector. A vaccine produced by AstraZeneca employs a Chimpanzee adenovirus vector and is approved for use in the European Union and other countries [4]. Since May 2022, 11 billion vaccine doses have been administered worldwide (https://covid19.who.int). However, SARS-CoV-2 variants have demonstrated immune escape in previously infected and fully vaccinated individuals. Compared to neutralization of alpha variant, serum from convalescent individuals was four-fold less effective against delta variant [5]. Similarly, serum from individuals who received two doses of Pfizer’s vaccine has neutralizing antibody titers against delta variant three- to five-fold lower than alpha variant. This immune escape was correlated to amino acid changes in the antigenic epitopes of the SARS-CoV-2 spike protein [5]. A study found that the omicron-neutralizing ability of serum from WA1-convalescent individuals was eight-fold lower than its WA1-neutralizing ability [6]. For individuals vaccinated with Pfizer, Moderna, J&J, or AstraZeneca vaccines, vaccine efficacy decreased by approximately 21 percentage points within one to six months after full vaccination, which was associated with waning immunity [7]. Due to waning immunity and the emergence of variants capable of immune escape, there is an urgent need for novel SARS-CoV-2 vaccine candidates with long-lasting protective immunity against the variants.

Parainfluenza virus 5 (PIV5) is a negative-sense, single-stranded, RNA virus in the family *Paramyxoviridae*. Its 15,246-nucleotide genome encodes for 8 proteins [8, 9]. Previously, recombinant PIV5 viruses expressing foreign genes from numerous pathogens, including influenza, rabies, respiratory syncytial virus, *Tuberculosis*, *Burkholderia*, and MERS-CoV have been generated and tested as vaccine candidates preclinically [10–15]. Because it actively replicates in the respiratory tract following intranasal immunization, PIV5-vectored vaccines can generate mucosal immunity that includes antigen-specific IgA antibodies and long-lived IgA plasma cells [12, 16]. Recently a PIV5-vectored vaccine expressing the spike protein from SARS-CoV-2 Wuhan (ancestral strain) (WA1; CVXGA1) has been shown to be efficacious in mice and ferrets [17]. A single, intranasal dose of CVXGA1 induced WA1-neutralizing antibodies and protected K18-hACE2 mice against lethal infection with SARS-CoV-2 WA1. Furthermore, a single, intranasal dose of CVXGA1 protected ferrets from SARS-CoV-2 WA1 infection and blocked contact transmission to cohoused naïve ferrets [17]. While these studies demonstrated its efficacy against SARS-CoV-2 WA1, CVXGA1’s efficacy against SARS-CoV-2 variants was not tested.

Golden Syrian hamsters have been proven susceptible to infection with SARS-CoV-2. Chan, et al. showed that following infection with WA1, hamsters lost weight for 6 days before starting to recover [18]. SARS-CoV-2 vRNA was detected in nasal turbinate, trachea, and lungs of infected hamsters, and peak infectious viral titer in the nasal turbinate and lungs was measured at 4 days post-infection [18]. While these studies used WA1 strain, several studies have since shown that hamsters are susceptible to infection with alpha and delta variants [19–21]. Among VOC alpha, delta, and omicron, delta causes the most weight loss and has the best viral replication in lungs of infected hamsters; omicron VOC, even with a high dose infection (2.5×10^6^ PFU per animal) did not cause weight loss, and replicates poorly in the lower respiratory tract of infected hamsters [22].

CVXGA1, recombinant PIV5 expressing S from SARS-CoV-2 WA1, is currently under phase 1 clinical trial in the US [23]. In this work, we examined efficacy of CVXGA1, and other recombinant PIV5 vaccines expressing S from SARS-CoV-2 beta, gamma, delta, or omicron, as a single-dose intranasal vaccine and as a boost following vaccination with two doses of COVID-19 mRNA vaccine against challenge infection with WA1, alpha, and delta in a Golden Syrian hamster model.

## Materials and Methods

### Cells

Vero E6 cells were maintained in Dulbecco’s modified Eagle media (DMEM) supplemented with 5% fetal bovine serum (FBS) plus 100 IU/mL penicillin and 100ug/mL streptomycin (1% P/S; Mediatech Inc, Manassas, VA, USA). Serum-free (SF) Vero cells were maintained in VP-SFM (ThermoFisher Scientific) plus 4mM GlutaMax (Gibco). Vero-TEMPRSS cells were obtained from Dr. Jeff Hogan, University of Georgia, and maintained in DMEM + 10% FBS + 1mg/mL G418. All cells were incubated at 37°C, 5% CO_2_.

### Plasmids and virus rescue

The construction of a plasmid encoding for PIV5 antigenome and generation of recombinant PIV5 were as previously described [24]. To construct plasmids encoding the antigenome of CVXGA1, CVXGA3, CVXGA5, CVXGA13, and CVXGA14, the Spike (S) genes from SARS-CoV-2 WA1, alpha, gamma, delta, and omicron, respectively, were placed as an additional open reading frame (ORF) transcription unit between the PIV5 SH and HN genes. The S cytoplasmic tail was replaced by the PIV5 fusion (F) protein cytoplasmic tail. To construct CVXGA2, encoding both the SARS-CoV-2 WA1 nucleoprotein (N) and S proteins, the N gene was placed as an additional ORF transcription unit between the PIV5 HN and L genes of CVXGA1. Primer sequences are available upon request. To generate recombinant PIV5 viruses CVXGA1, CVXGA2, CVXGA3, CVXGA5, CVXGA13, and CVXGA14, plasmids encoding the PIV5 antigenomic cDNA, the supporting plasmids (PIV5-NP, P, L), and T7 polymerase were transfected into serum-free (SF) Vero cells by FuGene transfection reagent (Fugent) or electroporation (Neon transfection system, Invitrogen). Recovered virus was amplified in SF Vero cells and the viral genomes were verified by RT-PCR and Sanger sequencing.

### Virus propagation

The recombinant PIV5 viruses were propagated in SF Vero cells at a multiplicity of infection (MOI) 0.001 PFU in VP-SFM + 4mM GlutaMax for 5 to 7 days at 37°C with 5% CO_2_. The media was collected and centrifuged at 1,500 rpm for 10 mins to pellet cell debris. The supernatant was mixed with 0.1 volume of 10X sucrose-phosphate-glutamate (SPG) buffer or 10X SPG + 10% Arginine, aliquoted, flash-frozen in liquid nitrogen, and stored at −80°C. The PIV5 virus stocks were titrated by plaque assay in Vero cells followed by immunostaining.

The SARS-CoV-2 viruses were propagated in Vero cells with DMEM + 1% FBS + 1X P/S. WA1 (BEI NR-52281) and alpha variant (USA/CA_CDC_5574/2020; BEI NR-54011) were obtained from BEI Resources. The omicron variant was provided by Dr. Jeff Hogan, University of Georgia. The delta variant was provided by Dr. Michael Gale, Jr., University of Washington. For isolation and production of delta variant stock, SARS-CoV-2 positive specimens with Ct < 33 were identified from reference testing [25]. The positive specimens were transferred to a biosafety level (BSL) 3 laboratory for virus culture. The virus transport medium (VTM) was first cleaned by filtering through Corning Costar Spin-X centrifuge tube filter (CLS8160), 0.1 mL of the cleaned VTM was used to infect Vero E6 cells ectopically expressing human ACE2 and TMPRSS2 (VeroE6-AT cells; a gift from Dr. Barney Graham, National Institutes of Health, Bethesda MD) in a 48-well plate. Two to four days post-infection when cytopathic effect, typical of SARS-CoV-2 infection, was observed, culture supernatants were collected and designated as a passage P0 virus stock. P1 virus stock cultures were grown in Vero E6/TMPRSS2 cells (JCRB1819) using P0 virus as inoculum. The titer of the P1 stock was measured by standard SARS-CoV-2 plaque assay as described [26].

P1 stock virus was verified with whole genome sequencing analysis. An aliquot of P1 stock was subject to RNA extraction (Zymo Research, R1040) and used as template to produce cDNA with SuperScript™ IV First-Strand Synthesis System (ThermoFisher, Waltham, MA, USA). The products were then subject to library production using the Swift SARS-CoV148 2 SNAP Version 2.0 kit (Swift Biosciences™, Ann Arbor, MI, USA) following the manufacturer’s instructions. Resulting libraries were quality-assessed using the Agilent 4200 TapeStation (Agilent Technologies, Santa155 Clara, CA, USA). Libraries with concentrations of 1.0 ng/μL were then sequenced on an Illumina NextSeq 500 (Illumina, San Diego, CA, USA) along with positive and negative controls.

The sequence data was processed through covid-swift-pipeline (https://github.com/greninger-lab/covid_swift_pipeline). Consensus genome sequences were generated by aligning the amplicon reads to the SARS-CoV-2 Wuhan-Hu-1 ancestral reference genome (NC_045512.2). For each genome, at least 1 million raw reads were acquired, representing >750x mean genome coverage and a minimum of 10x base coverage. Each consensus genome was then analyzed using the Phylogenetic Assignment of Named Global Outbreak Lineages (pangolin) tool to assign lineage based on the Pangolin dynamic lineage nomenclature scheme [27], defining the delta variant stock as Pangolin B.1.617.2.

### Immunofluorescence assay (IFA)

Immunofluorescence assays were performed to examine protein expression in virus-infected Vero cells. Vero cells were infected at MOI 0.01 with PIV5, CVXGA1, CVXGA2, CVXGA3, CVXGA5, CVXGA13, or CVXGA14 for 3 days before being fixed with 80% methanol. The cells were incubated with mouse anti-PIV5 V/P monoclonal antibody (PK 366), rabbit anti-SARS-CoV-2 S (Sino Biological catalog no. 40150-R007), or SARS-CoV-2 N (ProSci catalog no. 35-579) antibodies at 1:500 in PBS + 3% bovine serum albumin (BSA) for 1 hr. Next, the cells were washed with PBS and incubated with goat α-mouse Cy3 (KPL) or goat α-rabbit Cy3 (KPL) at 1:500 in PBS + 3% BSA for 30 mins. The cells were washed with PBS and imaged with an EVOS M5000 microscope (Thermo Fisher Scientific).

### Hamsters

Five-to-seven-week-old Golden Syrian hamsters were obtained from Charles River Laboratories. The hamsters were single housed in animal BSL2 (ABSL2) facilities with ad libitum access to food and water. Pre-challenge procedures were performed at the University of Georgia Biological Sciences Animal Facility. The hamsters were transferred to BSL3 facilities in the University of Georgia Animal Health Research Center (ABSL3) for the challenge and post-challenge procedures. The hamsters were anesthetized for immunization, blood collection, and challenge by intraperitoneal injection of 100 μL ketamine/acepromazine cocktail. All experiments were performed in accordance with protocols approved by the Institutional Animal Care and Use Committee at the University of Georgia.

### Immunization and challenge of hamsters

To administer intranasal immunizations, anesthetized hamsters were placed on their backs, a pipette was used to dispense 100 μL inoculum onto their noses, and the inoculum was allowed to drain into their respiratory tracts. They were recovered on heating pads.

A COVID-19 mRNA vaccine was obtained from a clinical site, reconstituted to 200 μg/mL, aliquoted, and stored at −80°C. Two μg mRNA vaccine in 50 μL was administered via intramuscular injection.

For study AE19 (Figure 4), hamsters (n=8) received a single intranasal immunization of 100 μL of 10^5^ plaque-forming units (PFU) PIV5, CVXGA1, CVXGA2, CVXGA3, or CVXGA5. At 28 days post-immunization (dpi), blood was collected from the hamster saphenous vein for serological analysis. At 36 dpi, the hamsters were anesthetized: four hamsters were challenged intranasally with 30 μL 10^3^ PFU of SARS-CoV-2 Wuhan strain (WA1), and the remaining four hamsters were challenged with 10^3^ PFU SARS-CoV-2 alpha variant (CA; BEI NR54011) as previously reported by Blanchard, et al. [28]. Following challenge infection, the hamster weights were monitored for 5 days. At 5 days post-challenge (dpc), the hamsters were euthanized, and the hamster lungs were harvested, resuspended in 2 mL DMEM + 2% FBS + 1X antibiotic/antimycotic, homogenized, aliquoted, and stored at −80°C. SARS-CoV-2 viral burden in lung homogenate was quantified via plaque assay and real-time quantitative reverse transcription polymerase chain reaction (RT-qPCR).

For study AE23 (Figure 5), hamsters received intramuscular (i.m.) immunizations of 100 μL PBS (n=20, group 1) or 2 μg mRNA COVID-19 vaccine (n=20, group 2). At 21 dpi, hamsters that received the mRNA vaccine were boosted with the mRNA vaccine. At 28 dpi, blood was collected from the hamster saphenous vein for serological analysis. At 35 dpi following initial immunization, hamsters that received PBS during the first immunization received either 100 μL of PBS intranasally (i.n.) (n=5, group 1A), 3×10^5^ PFU CVXGA1 (n=5, group 1B), 2×10^5^ PFU CVXGA3 (n=5, group 1C), or 1.5×10^5^ PFU CVXGA13 (n=5, group 1D). Group 2 hamsters that received two doses of mRNA received 100 μL of PBS (n=4, group 2A), 3×10^5^ PFU CVXGA1 (n=4, group 2B), 2×10^5^ PFU CVXGA3 (n=4, group 2C), 1.5×10^5^ PFU CVXGA13 (n=4, group 2D) i.n., or a third dose of mRNA i.m. (n=4, group 2E). Hamsters were anesthetized for intranasal immunizations but not intramuscular injections. Blood was collected at 54 dpi. At 63 dpi, the hamsters were challenged with 10^4^ PFU SARS-CoV-2 delta variant. Following challenge infection, hamster weights were monitored for 5 days. At 5 dpc, the hamsters were euthanized, their lungs were harvested, and the SARS-CoV-2 viral burden was quantified via plaque assay and RT-qPCR.

For study AE24 (Figures 6 and 7), hamsters received 100 μL of PBS i.n. (n=5, Group 1), 2 μg mRNA COVID-19 vaccine i.m. (n=25, Group 2), or 100 μL 7×10^4^ PFU CVXGA1 i.n. (n=10, Groups 3 & 4). At 29 dpi, the hamsters that received the mRNA vaccine were boosted with the mRNA vaccine i.m. and the group 3 hamsters received another dose of CVXGA1 i.n. At 91 dpi following initial immunization, hamsters who received two doses of mRNA received either 2 μg mRNA vaccine i.m. (n=5, Group 2A) or 100 μL of 7×10^4^ PFU CVXGA1 (n=5, Group 2B), 10^3^ PFU CVXGA13 (n=5, Group 2C), PBS i.n. (n=5, Group 2D), or 10^4^ PFU CVXGA14 (n=5, Group 2E). Hamsters were anesthetized for intranasal immunizations but not intramuscular injections. At 36 and 108 dpi, blood was collected via the hamster gingival vein. At 116 dpi, the hamsters were challenged with 30 μL of 10^4^ PFU SARS-CoV-2 delta variant. Following challenge infection, hamster weights were monitored for 5 days. The hamster lungs were harvested, and SARS-CoV-2 viral burden was quantified via plaque assay and RT-qPCR.

All animal experiments were performed according to the protocols approved by the Institutional Animal Care and Use Committee at the University of Georgia.

### Enzyme-linked immunosorbent assay (ELISA)

To quantify the anti-SARS-CoV-2 S and RBD humoral response, hamster serum was analyzed via ELISA. Immulon® 2HB 96-well microtiter plates were coated with 100 μL SARS-CoV-2 S or RBD at 1 μg/mL. For all ELISAs, plates were coated with SARS-CoV-2 S and RBD from the WA1 strain, which were produced and purified as described previously [17]. Hamster serum was serially diluted two-fold and incubated on the plates for 2 hrs. Horseradish peroxidase-labelled goat anti-mouse IgG secondary antibody (Southern Biotech, Birmingham, Alabama) was diluted 1:2000 and incubated on the wells for 1 hr. The plates were developed with KPL SureBlue Reserve TMB Microwell Peroxidase Substrate (SeraCare Life Sciences, Inc., Milford, Massachusetts), and OD_450_ values were obtained with a BioTek Epoch Microplate Spectrophotometer (BioTek, Winooski, Vermont). Antibody titers were calculated as log_10_ of the highest serum dilution at which the OD_450_ was greater than two standard deviations above the mean OD_450_ of naïve serum.

### Neutralization assays

To quantify the SARS-CoV-2-neutralizing antibodies generated by the hamsters, microneutralization assays were performed in a BSL 3 facility. Hamster serum was heat-inactivated at 56°C for 45 mins and serially diluted two-fold. The serum was mixed 1:1 with 6×10^3^ focus-forming units (FFU)/mL SARS-CoV-2 WA1, delta, or omicron variants. The serum/virus mixture was incubated at 37°C for 1 hr before being incubated on 96-wells of Vero cells for WA1 or Vero TEMPRSS2 cells for delta and omicron, respectively. One hour post-infection, a methylcellulose overlay (DMEM + 5% FBS + 1% P/S + 1% methylcellulose) was added on top of the serum/virus mixture. The plates were incubated at 37°C, 5% CO_2_ for 24 hrs. After removal of the methylcellulose overlay, the wells were washed with PBS, and the cells were fixed with 60% methanol/40% acetone, followed by immunostaining with anti-SARS-CoV-2 N antibody (ProSci catalog no. 35-579). The number of infected cells were quantified via Cytation 7 imaging reader (BioTek). Neutralization titers were calculated as log_10_ of the highest serum dilution at which the virus infectivity was reduced by at least 50%.

### Plaque assay for infectious virus titer

To quantify infectious SARS-CoV-2 in lung homogenates, plaque assays were performed. For the plaque assays, lung homogenates were serially diluted in DMEM + 2% FBS + 1% antibiotic/antimycotic and added to 12-well plates of Vero E6 cells for SARS-CoV-2 WA1 and alpha variant or Vero TEMPRSS2 cells for delta and omicron variants. At 1 hour post-infection, the inoculum was removed, and a methylcellulose overlay (500mL Opti-MEM + 0.8% methylcellulose + 2% FBS + 1% antibiotic/antimycotic) was added to the wells. Following incubation for 3 days, the overlay was removed, and the cells were fixed with 60% methanol/40% acetone. After staining with crystal violet, the number of plaques were counted, with viral titers expressed as PFU/mL of lung homogenate.

### qPCR

SARS-CoV-2 viral RNA levels were quantified by RT-qPCR. SARS-CoV-2 virus was inactivated by mixing 100 μL lung homogenate with 900 μL TRIzol (Invitrogen). Using a QIAgen RNA extraction kit, RNA was extracted from 140 μL homogenate/TRIzol and eluted in 15 μL of elution buffer, of which 5 μL was used in the qRT-PCR reaction. qRT-PCR was performed according to the protocol described in the “CDC 2019-Novel Coronavirus (2019-nCoV) Real-Time RT-PCR Diagnostic Panel…Instructions for Use” (page 26; https://www.fda.gov/media/134922/download) with Applied Biosystems TaqPath One Step RT qPCR Master Mix and SARS-CoV-2 Research Use Only qPCR Primer and Probe Kit primer/probe mix N1. SARS-CoV-2 viral RNA was extracted from 140 μL of SARS-CoV-2 WA1, alpha variant, and delta variant viruses of known titers and eluted in 15 μL of elution buffer. The viral RNA was serially diluted 10-fold and 5 μL from each dilution was used in the RT-qPCR assay. To generate a standard curve, the viral titer was plotted on the x-axis and the CT value was plotted on the y-axis. The standard curves were used to calculate the CT value that corresponds to 1 PFU/rxn in virus stock and hamster lung homogenates. The CT value of RNA extracted from sterile elution buffer was designated the PCR negative cutoff.

## Results

### Construction and characterization of PIV5-vectored SARS-CoV-2 vaccines

We previously generated a PIV5-vectored vaccine for SARS-CoV-2 by inserting the SARS-CoV-2 WA1 S gene, which had the cytoplasmic tail of the S protein replaced with the cytoplasmic tail from the PIV5 F protein, between the PIV5 SH and HN genes (CVXGA1). We showed that a single, intranasal dose of CVXGA1 protects K18-hACE2 mice from lethal infection with the WA1 strain, the initial circulating strain in the US, and blocks contact transmission in ferrets [17]. To determine whether expressing the N protein of SARS-CoV-2 as an additional antigen enhances protection afforded by the S antigen alone, we generated PIV5 expressing both S and N (CVXGA2). During our study period, SARS-CoV-2 VOCs emerged and some became dominant strains at different times. Thus, we generated PIV5-vectored vaccine candidates expressing S from SARS-CoV-2 VOC in a similar manner as CVXGA1 (Figure1): variants beta (CVXGA3), gamma (CVXGA5), delta (CVXGA13), and omicron (CVXGA14) (collectively called CVXGA vaccines). All variant S genes had the cytoplasmic tail replaced with the PIV5 F cytoplasmic tail.

The vaccine viruses were recovered as previously described, and their genomes were confirmed by RT-PCR and sequencing [29]. Compared to PIV5 vector-infected cells, all PIV5-vectored vaccines had increased syncytia, indicating that SARS-CoV-2 S is functional (Figure 2A).

**Figure 1.**
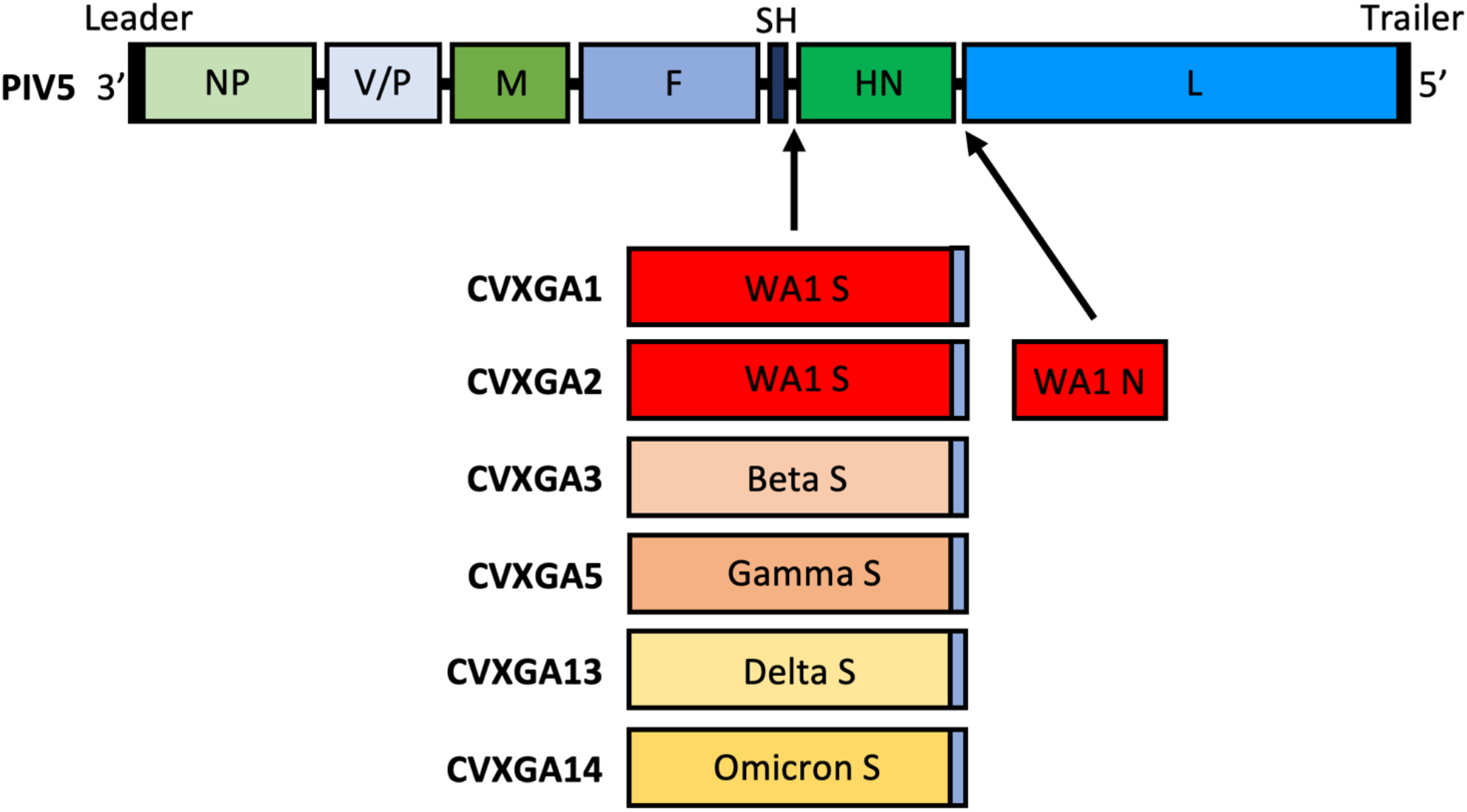
Schematics of PIV5 and CVXGA vaccines. The PIV5 genome has 7 genes 3’ to 5’: NP, V/P, M, F, SH, HN, L. The S genes from SARS-CoV-2 WA1 (CVXGA1), beta variant (CVXGA3), gamma variant (CVXGA5), delta variant (CVXGA13), and omicron variant (CVXGA14) had their cytoplasmic tails replaced with the PIV5 F cytoplasmic tail and inserted between PIV5 SH and HN genes. CVXGA2 also has SARS-CoV-2 WA1 N inserted between PIV5 HN and L genes.

**Figure 2.**
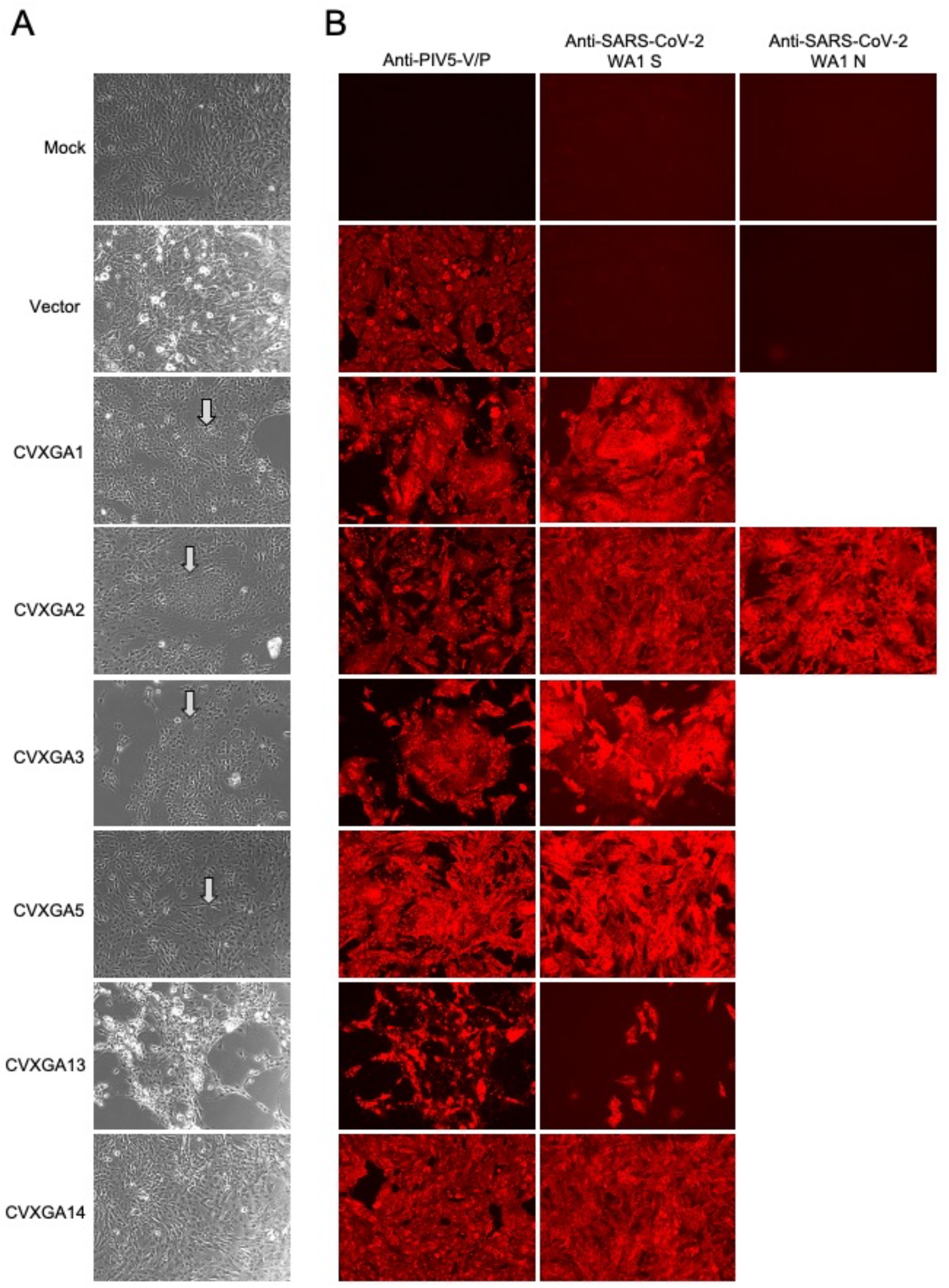
CVXGA vaccine antigen expression. Vero cells were infected at MOI 0.01 for 3 days. (A) Cell-to-cell fusion induced by SARS-CoV-2 S expression was imaged at 10X with an Evos M5000 microscope. Arrows indicate syncytium, multinucleated cells. (B) Intracellular expression of PIV5-V/P, SARS-CoV-2-S, and SARS-CoV-2-N was detected by anti-PIV5 V/P, -SARS-CoV-2 S, or SARS-CoV-2 N antibodies, followed by Cy3-conjugated secondary antibody, and imaged at 10X with an EVOS M5000 microscope (Thermo Fisher Scientific).

To further confirm antigen expression, Vero cells were infected at MOI 0.01 with PIV5, CVXGA1, CVXGA2, CVXGA3, CVXGA5, CVXGA13, or CVXGA14 and assayed for immunofluorescence with WA1 S-specific antibody or N-specific antibody for CVXGA2. As expected, S expression was detected in cells infected with CVXGA1, 2, 3, 5, 13 and 14. Additionally, SARS-CoV-2 N expression was detected in cells infected with CVXGA2 (Figure 2B).

### PIV5-vectored SARS-CoV-2 vaccines induce an anti-S humoral response in hamsters

To test efficacy of PIV5-vectored COVID-19 vaccine candidates in hamsters, their ability to induce S-specific antibody responses in hamsters was examined. We immunized Golden Syrian hamsters with a single, intranasal dose of 10^5^ plaque-forming units (PFU) PIV5 vector, CVXGA1, CVXGA2, CVXGA3, or 5×10^2^ PFU CVXGA5 (Figure 3A). While hamsters immunized with PIV5 vector had no detectable anti-SARS-CoV-2-S binding antibodies at day 28 dpi, a single intranasal dose of CVXGA1, CVXGA2, or CVXGA3 induced mean ELISA antibody titers of over 10,000. Even CVXGA5, at a lower immunization dose, was able to induce an anti-S ELISA titer greater than 9,333 (Figure 3B).

**Figure 3.**
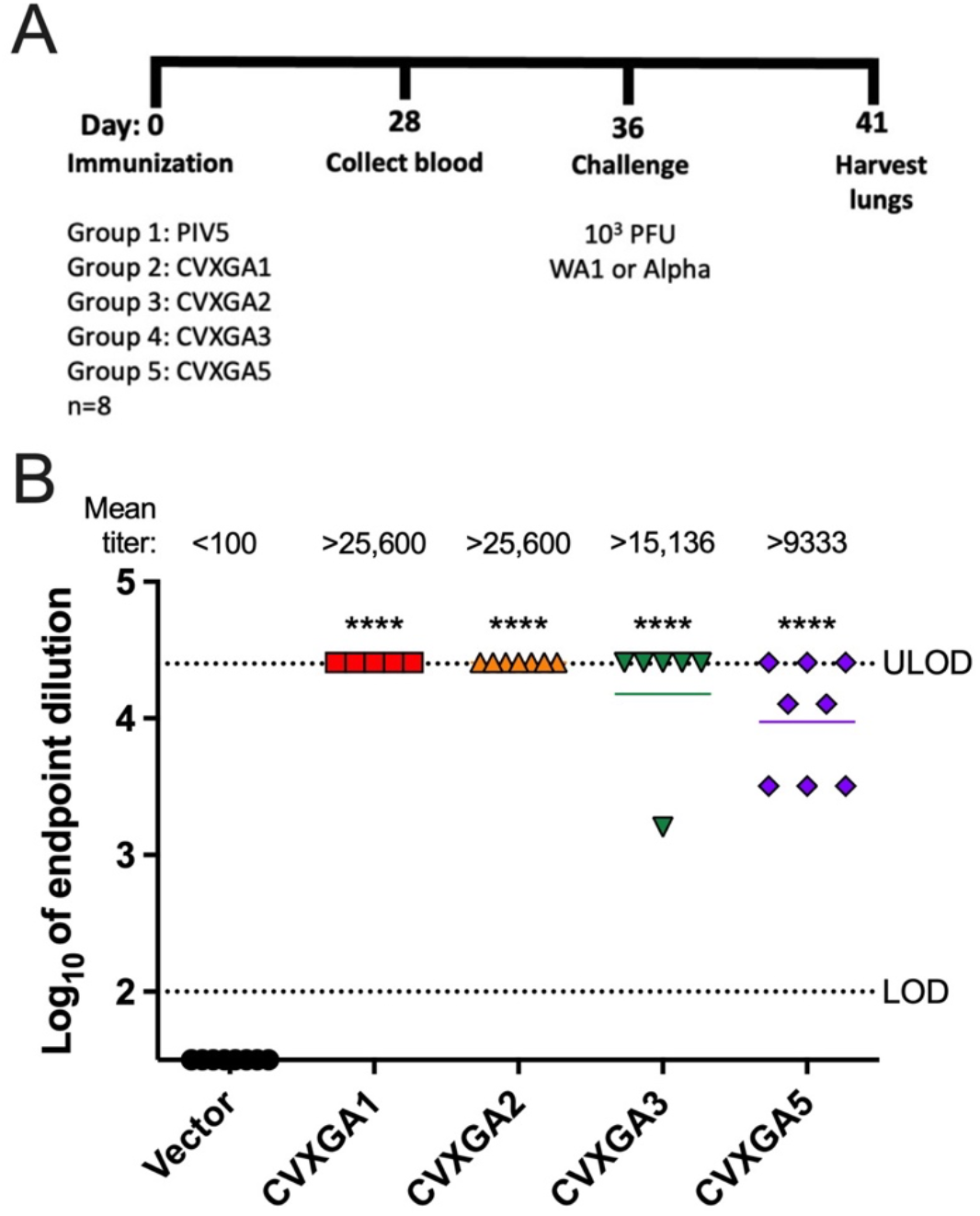
Immunization of hamsters with CVXGA1, CVXGA2, CVXGA3, and CVXGA5 induces anti-SARS-CoV-2 S IgG antibodies. (A) Schematic of hamster study AE19 immunization. Golden Syrian hamsters (n=8) were intranasally immunized with 100 μL of 10^5^ PFU PIV5, CVXGA1, CVXGA2, CVXGA3, or CVXGA5. Blood was collected at 28 dpi. At 36 dpi, four hamsters from each group were challenged with 10^3^ PFU of SARS-CoV-2 Wuhan strain (WA1) and the remaining four hamsters were challenged with 10^3^ PFU SARS-CoV-2 alpha variant. Following challenge infection, the hamster weights were monitored daily before terminating the study at 5 dpc to collect lung tissues. (B) Anti-SARS-CoV-2 S IgG antibody titers were quantified by ELISA. Antibody titers were calculated as log_10_ of the highest serum dilution at which the OD_450_ was greater than two standard deviations above the mean OD_450_ of naïve serum. The lower limit of detection (LOD) and upper limit of detection (ULOD) are indicated by the dotted lines. Bars represent the geometric means. Comparing each group to the vector control, statistical significance was calculated with one-way ANOVA (**** p < 0.0001).

### CVXGA vaccines protect against homologous and heterologous challenges

To assess the efficacy of PIV5-vectored SARS-CoV-2 vaccines against homologous and heterologous virus challenges, CVXGA-immunized hamsters were challenged with either 10^3^ PFU SARS-CoV-2 WA1 (USA-WA01/2020) or alpha variant (CA; BEI NR54011) at 36 dpi. The hamster weights were monitored daily for 5 days post-challenge (dpc). Following challenge with WA1, hamsters immunized with PIV5 vector lost weight and did not recover before the study was terminated. In contrast, hamsters immunized with CVXGA1, CVXGA2, CVXGA3, or CVXGA5 lost weight at day 1 post-challenge but returned to pre-challenge weights 3 to 4 dpc (Figure 4A). While challenge with alpha variant induced less severe weight loss in PIV5 vector-immunized hamsters, hamsters immunized with CVXGA1, CVXGA2, CVXGA3, or CVXGA5 had significantly higher body weights compared to PIV5 vector-immunized hamsters at 5 dpc. In both virus challenge groups, hamsters immunized with CVXGA2 had the highest mean body weight gains following challenge (Figure 4B), suggesting that the SARS-CoV-2 N antigen provided additional protection.

**Figure 4.**
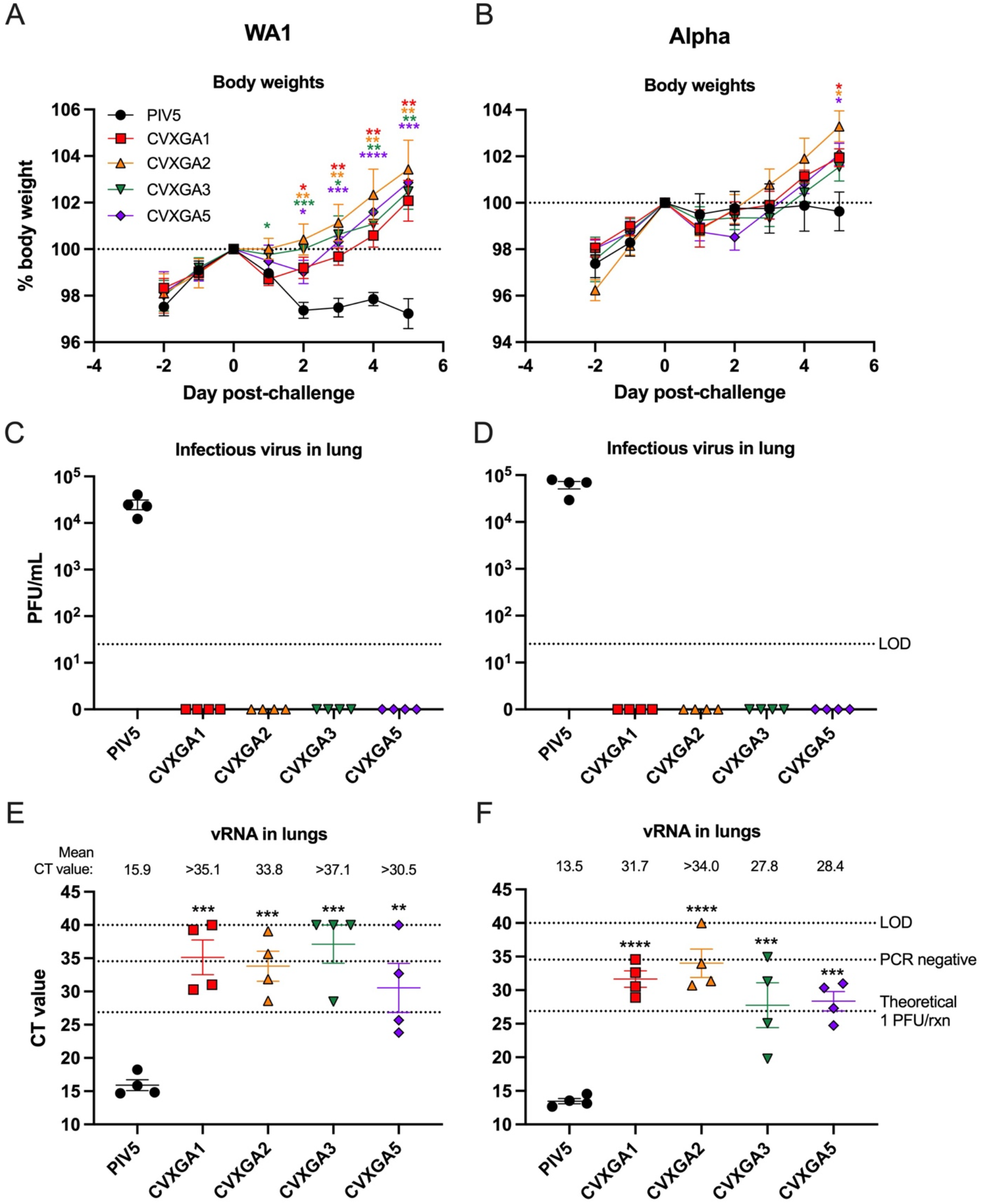
Immunization with CVXGA1 protects hamsters from challenge with SARS-CoV-2 WA1 and alpha variant. Following challenge with WA1 (A) or alpha variant (B), hamster weights were monitored daily for five days and graphed as percent day 0 weight. Statistical significance was calculated for each timepoint between each group and PIV5-immunized hamsters with t tests (* p ≤ 0.05, ** p < 0.01, *** p < 0.001, **** p < 0.0001). At 5 dpc with WA1 (C) or alpha variant (D), viral load in hamster lung was quantified via plaque assay in Vero E6 cells and graphed as PFU/mL lung homogenate. The limit of detection (LOD) is indicated by the dotted line. Error bars represent the standard error of the means. SARS-CoV-2 WA1 (E) or alpha variant (F) vRNA load in lung homogenates was quantified via RT-qPCR. The cycle threshold (Ct) value for each sample is presented and error bars represent the standard error of the means. The known viral titers of WA1 and alpha variant were used to generate standard curves for E and F, respectively, and the Ct values equating to 1 PFU per reaction (rxn) were calculated. The Ct value generated from RNA extracted from sterile water is denoted by a dotted line labelled PCR negative. The limit of detection (LOD) is indicated by a dotted line at Ct value = 40, the number of PCR cycles. Comparing each group to PIV5-immunized hamsters, statistical significance was calculated with one-way ANOVA (** p < 0.01, *** p < 0.001).

To quantify challenge viral burden in the lungs, infectious virus in lung homogenates was quantified by plaque assay. Hamsters immunized with PIV5 vector had infectious SARS-CoV-2 virus titer greater than 4 log_10_ PFU/mL lung homogenate following challenge with WA1 or alpha variant, while no infectious WA1 or alpha variant was detected in hamsters immunized with CVXGA1, CVXGA2, CVXGA3, or CVXGA5 (Figures 4C & D; limit of detection, LOD, 25 PFU/mL). Viral RNA in the lung was quantified by RT-qPCR. Hamsters immunized with PIV5 vector and challenged with WA1 or alpha variant had mean cycle threshold (CT) values of 15.9 and 13.5, respectively (Figures 4E & F). Following challenge with WA1 or alpha variant, hamsters that received a single intranasal dose of CVXGA1 or CVXGA2 had CT values indicative of less than 1 PFU per reaction (PFU/rxn). CVXGA3-immunized hamsters had CT values indicative of less than 1 PFU/rxn following challenge with WA1 but two hamsters had CT values equating to 1 or 96 PFU/rxn following challenge with alpha variant (Figures 4E & F, Supplemental figure 1). A single dose of CVXGA2 performed the best against heterologous challenge with alpha variant (Figure 4F), suggesting that SARS-CoV-2 N might offer additional protection.

### CVXGA vaccines protect against delta challenge

As of May 2022, delta variant is one of two VOCs circulating in the United States (https://www.who.int/activities/tracking-SARS-CoV-2-variants). Therefore, we assessed the efficacy of our lead vaccine candidate, CVXGA1, against heterologous challenge with delta variant and tested a vaccine expressing S from delta variant against homologous challenge. Hamsters received a single, intranasal dose of PBS or 10^5^ PFU of CVXGA1, CVXGA3, or CVXGA13. At 19 dpi, blood was collected and serum anti-SARS-CoV-2 WA1 S and -RBD IgG antibodies were quantified via ELISA (Figure 5A). CVXGA1, CVXGA3, or CVXGA13 elicited anti-S antibodies with mean titers of over 10,000 (Figure 5B). Interestingly, hamsters immunized with CVXGA13 had the highest level of anti-WA1 RBD antibodies with a mean titer of 50,119 (Figure 5C).

**Figure 5.**
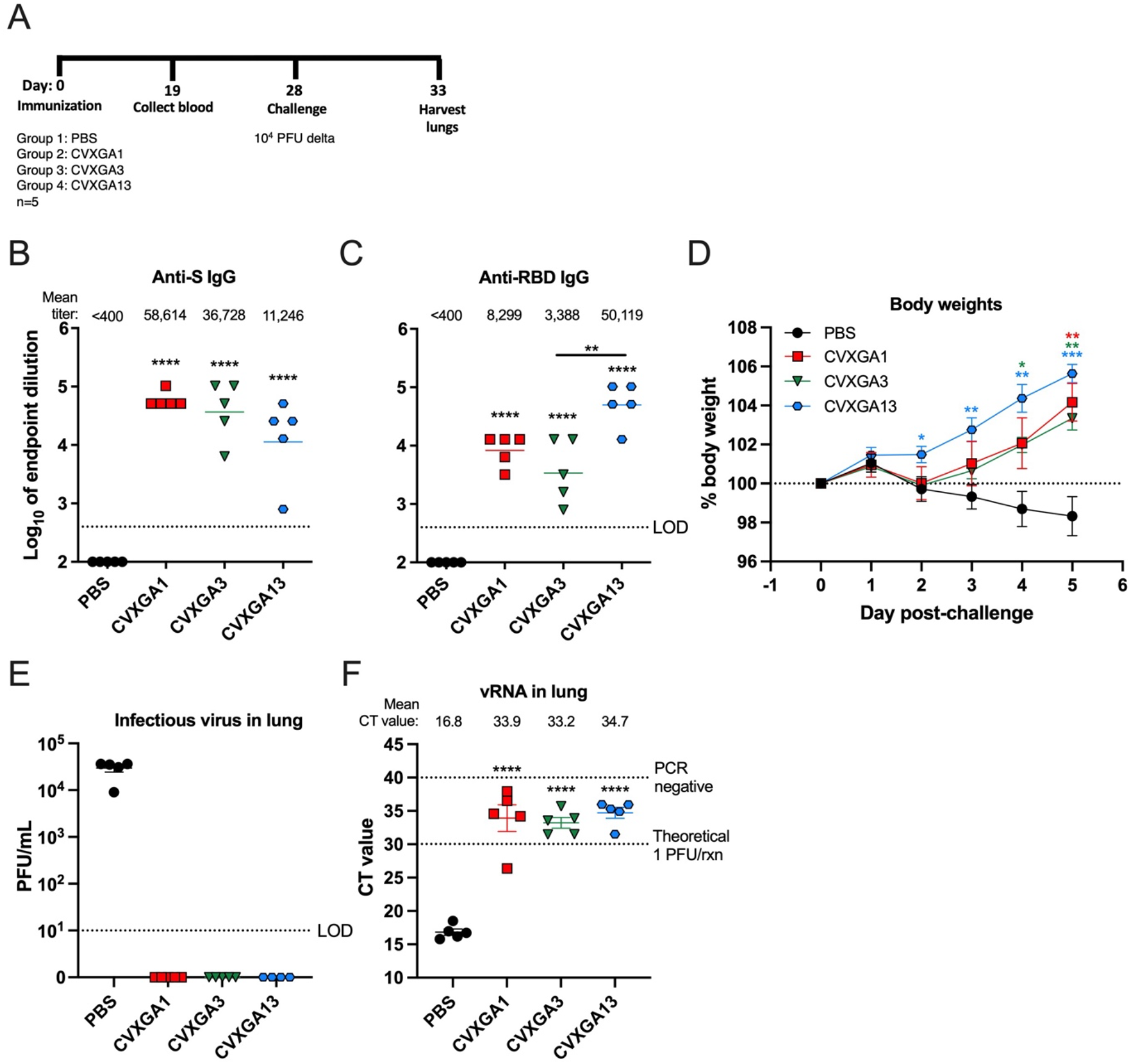
Immunization with CVXGA1 protects hamsters from challenge with SARS-CoV-2 delta variant. (A) Schematic of hamster study AE23 immunization. Golden Syrian hamsters (n=5) were intranasally immunized with 100 μL PBS, 3×10^5^ PFU CVXGA1, 2×10^5^ PFU CVXGA3, or 1.5×10^5^ PFU CVXGA13. Blood was collected at 19 dpi. At 28 dpi, the hamsters were challenged with 10^4^ PFU of SARS-CoV-2 delta variant. Following challenge infection, the hamster weights were monitored daily before terminating the study and harvesting lungs at 5 dpc. Anti-SARS-CoV-2 WA1 S (B) and RBD (C) IgG antibodies were quantified via ELISA at 19 dpi. Antibody titer was calculated as log_10_ of the highest serum dilution at which the OD_450_ was greater than two standard deviations above the mean OD_450_ of naïve serum. The limit of detection (LOD) is indicated by the dotted line. Error bars represent the standard error of the means. Comparing each group to the vector control, statistical significance was calculated with one-way ANOVA (** p < 0.01, **** p < 0.0001). (D) Following challenge infection, hamster body weights were monitored daily for five days and graphed as percent of day 0 weight. Statistical significance was calculated for each timepoint between each group and PIV5-immunized hamsters with t tests (* p ≤ 0.05, ** p < 0.01, *** p < 0.001). (E) Viral load in lung homogenate was quantified via plaque assay in Vero TEMPRSS cells and graphed as PFU/mL lung homogenate. The limit of detection (LOD) is indicated by the dotted line. Error bars represent the standard error of the means. (F) RNA was extracted from lung homogenate and SARS-CoV-2 delta vRNA was quantified via RT-qPCR. The cycle threshold (Ct) value for each sample is presented and error bars represent the standard error of the means. The known viral titer of delta variant was used to generate a standard curve to calculate the Ct value equating to 1 PFU per reaction (rxn). The limit of detection (LOD), PCR negative, is indicated by a dotted line at Ct value = 40, the number of PCR cycles.

At 28 dpi, the hamsters were intranasally challenged with 10^4^ PFU SARS-CoV-2 delta variant and their weights were monitored for 5 days. Beginning at 2 dpc, hamsters immunized with PBS experienced weight loss that steadily declined until the study was terminated. In contrast, hamsters immunized with CVXGA1, CVXGA3, or CVXGA13 experienced weight gain after 2 dpc. While not statistically significant, hamsters immunized with CVXGA13 had greater weight gain than hamsters immunized with CVXGA1 or CVXGA3, indicating that PIV5 expressing S from SARS-CoV-2 delta variant protected hamsters the best from weight loss following homologous challenge with delta variant (Figure 5D). To further assess the vaccine efficacy, challenge virus load in lungs at 5 dpc were examined by plaque assay and RT-qPCR. While hamsters immunized with CVXGA1, CVXGA3, or CVXGA13 did not have detectable infectious challenge virus in their lung homogenates, hamsters immunized with PBS had a mean titer of 3 x 10^4^ PFU/mL lung homogenate (Figure 5E). Four of five hamsters immunized with CVXGA1 and all hamsters immunized with CVXGA3 or CVXGA13 had CT values indicative of no infectious virus. One CVXGA1-immunized hamster had a CT value of 26.39 (Figure 5F). The weight loss and lung viral burden data from this study demonstrated that single, intranasal doses of CVXGA1 and CVXGA3 protect hamsters from heterologous challenge and CVXGA13 protects hamsters from homologous challenge with delta variant. The protective effect offered by CVXGA13 against homologous delta virus is the best among the three CVXGA vaccine candidates.

### CVXGA1 generates longer lasting immunity

As of May, 2022, 67 percent of individuals in the United States are fully vaccinated against COVID-19 (https://ourworldindata.org/covid-vaccinations?country=USA). However, vaccine-induced immunity wanes over time, making vaccines less effective against SARS-CoV-2 variants [5–7]. To assess the longevity of CVXGA1, we compared one (1X CVXGA1) and two (2X CVXGA1) intranasal doses of CVXGA1 to two intramuscular doses of a mRNA COVID-19 vaccine (2X mRNA) in hamsters over time (Figure 6A). Blood was collected 7 and 79 days following the second immunization and anti-SARS-CoV-2-S IgG antibodies were quantified via ELISA. Hamsters who received 2X mRNA had highest anti-S ELISA titer, and 2X CVXGA1-immunized hamsters had higher anti-S titers than 1X CVXGA1-immunized hamsters, whose mean anti-WA1 S antibody titer was 50,699 on day 36 but the titers on day 108 are comparable for all three vaccines (Figure 6B). To assess the neutralizing ability of the antibodies, the hamster serum was tested in microneutralization assays with SARS-CoV-2 WA1, delta variant, and omicron variant. While 2X mRNA generated a high level of anti-WA1 neutralizing antibody (4,257 at 7 days post-second dose), 2X CVXGA1 generated comparable levels of anti-S ELISA titers but higher levels of anti-WA1 neutralizing antibodies at day 108 (Figure 6B & C). Consistent with reduced cross-reactivity of mRNA-generated antibodies with delta and omicron variants, delta- and omicron-neutralizing antibody levels were lower than anti-WA1 in hamsters immunized with 2X mRNA at 531 and 286 respectively at 7 days post-second dose. As expected, 1X and 2X CVXGA1 generated anti-delta neutralization levels lower than anti-WA1 neutralization levels and even lower for anti-omicron (Figure 6C).

**Figure 6.**
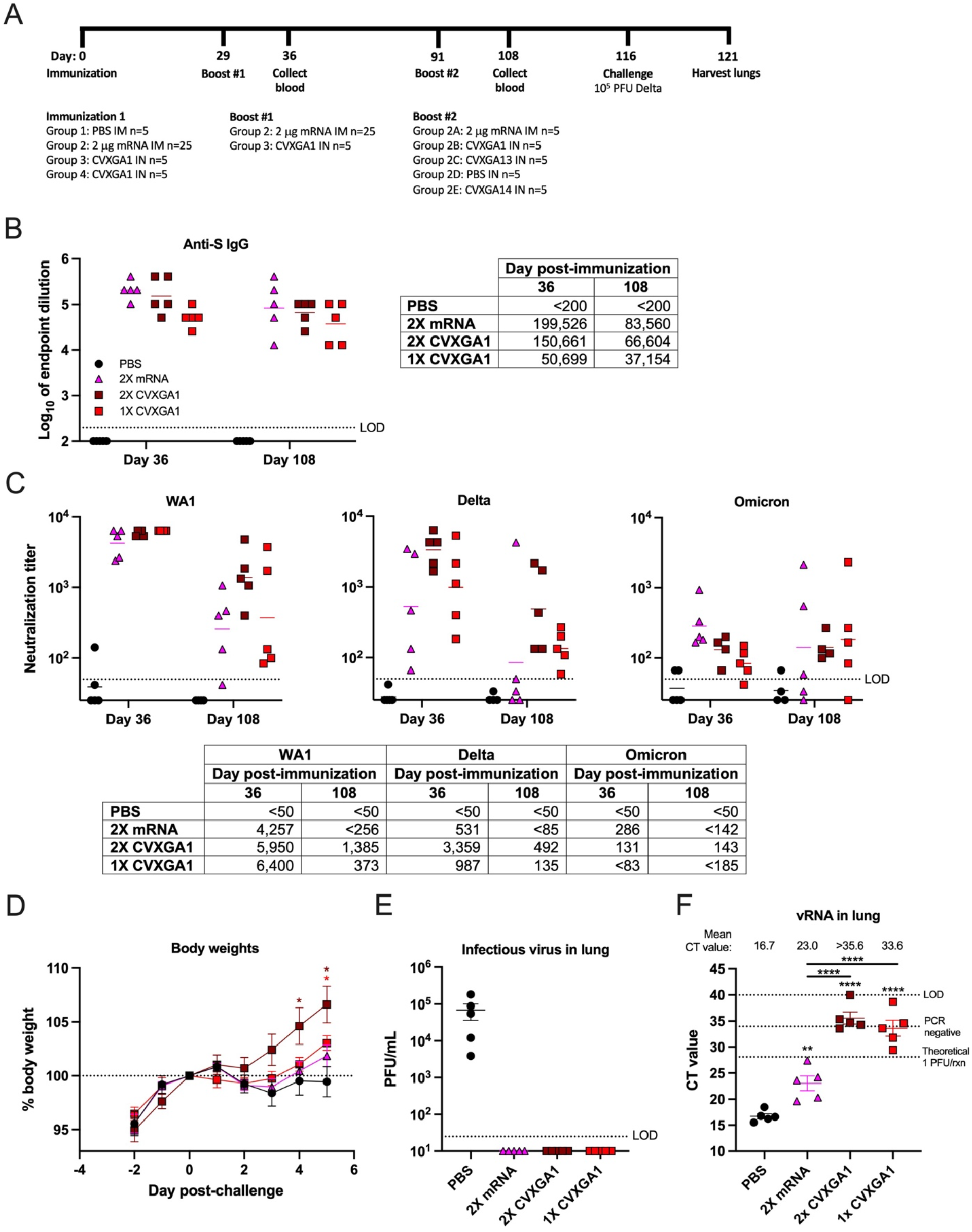
Immunization with a COVID-19 mRNA vaccine, one dose of CVXGA1, or two doses of CVXGA1 protects hamsters against challenge with delta variant. (A) Schematic of hamster study AE24 immunization. Golden Syrian hamsters received 100 μL PBS intranasally (n=5, Group 1), 2 μg mRNA COVID vaccine (n=25, Group 2), or 100 μL 7×10^4^ PFU CVXGA1 (n=10, Groups 3 & 4). At 29 dpi, hamsters that received the mRNA vaccine were boosted with the mRNA vaccine and group 3 hamsters received another dose of CVXGA1. At 91 dpi following initial immunization, hamsters who received two doses of mRNA received 2 μg mRNA vaccine (n=5, Group 2A), 7×10^4^ PFU CVXGA1 (n=5, Group 2B), 10^3^ PFU CVXGA13 (n=5, Group 2C), PBS i.n. (n=5, Group 2D), or 10^4^ PFU CVXGA14 (n=5, Group 2E). Blood was collected at 36 and 108 dpi. At 116 dpi, the hamsters were challenged with 10^5^ PFU SARS-CoV-2 delta variant. Following challenge infection, hamster weights were monitored for 5 days and the lungs were harvested. (B) Anti-SARS-CoV-2 WA1 S IgG antibodies were quantified via ELISA at 36 and 108 days post-immunization. Antibody titer was calculated as log_10_ of the highest serum dilution at which the OD_450_ was greater than two standard deviations above the mean OD_450_ of naïve serum. The limit of detection (LOD) is indicated by the dotted line. The geometric means are presented in the table and represented as bars on the graph. (C) Microneutralizing antibody titers against SARS-CoV-2 WA1, delta, or omicron were calculated as log_10_ of the highest serum dilution at which the virus infectivity was reduced by at least 50%. The limit of detection (LOD) is indicated by the dotted line. The geometric means are presented in the table and represented as bars on the graph. (D) Following challenge, hamster weights were monitored daily for five days and graphed as percent of day 0 weight. Statistical significance was calculated for each timepoint between each group and PBS-immunized hamsters with t tests (* p ≤ 0.05). (E) Viral load in lung homogenate at 5 dpc was quantified via plaque assay in Vero TEMPRSS cells and graphed as PFU/mL lung homogenate. The limit of detection (LOD) is indicated by the dotted line. Error bars represent the standard error of the means. (F) Viral RNA load in lung homogenate was quantified via RT-qPCR. The cycle threshold (Ct) value for each sample is presented and error bars represent the standard error of the means. The known viral titer of delta variant was used to generate a standard curve and calculate the Ct value equating to 1 PFU per reaction (rxn). The Ct value generated from RNA extracted from sterile water is denoted by a dotted line labelled PCR negative. The limit of detection (LOD) is indicated by a dotted line at Ct value = 40, the number of PCR cycles. Error bars represent the standard error of the means. Statistical significance was calculated with one-way ANOVA (** p < 0.01, **** p < 0.0001).

To compare longevity of antibody responses in the hamsters, sera were collected 79 days after boost (day 108 after initial immunization). Anti-S ELISA titers dropped during this time to 41.9%, 44.2%, and 73.3% for 2X mRNA, 2X CVXGA1, and 1X CVXGA1, respectively (Figure 6B). Reduction of neutralizing antibody titers in 2X mRNA-immunized hamsters was substantial with 20, 60, and 40 percent of hamsters having no detectable WA1-, delta-, and omicron-neutralizing antibodies at 79 days post-boost. In contrast, serum from all hamsters immunized with 2X CVXGA1 maintained levels of neutralizing antibodies against WA1, delta variant, and omicron variant better than the 2X mRNA vaccine group (Figure 6C).

Eighty-seven days after the second immunization, the hamsters were challenged with delta variant and the hamster weights were monitored for 5 days. Compared to hamsters who were immunized with PBS, hamsters who received two doses of CVXGA1 had significant weight gain following challenge. Hamsters who received two doses of mRNA vaccine did experience weight gain but it was not statistically significant from the PBS group (Figure 6D). Viral burden in the hamster lungs at 5 dpc was tested by plaque assay and RT-qPCR. While PBS-immunized hamsters had mean lung titers of 5.2 x 10^3^ FFU/mL lung homogenate, none of the vaccinated hamsters had detectable infectious virus in their lung homogenate (Figure 6E). However, all hamsters immunized with two doses of mRNA vaccine had SARS-CoV-2 vRNA levels indicative of infectious virus with the mean CT value equating to 38 PFU per RT-qPCR reaction (Figure 6F).

### Boosting with CVXGA improves protection of hamsters immunized with mRNA COVID-19 vaccine

Due to large populations having already been immunized with COVID-19 vaccines, we investigated the use of CVXGA vaccines as a booster in the hamster model (Figure 6A, group 2). We used 2X mRNA immunization as a starting point for comparison. Hamsters were first immunized with two doses of mRNA COVID-19 vaccine. At 62 days after second dose of mRNA vaccination, hamsters were boosted with mRNA (total 3X mRNA vaccine doses), CVXGA1, CVXGA3, CVXGA14, or PBS. Seventeen days after the third immunization, blood was collected from the hamsters and anti-WA1 S IgG antibodies were quantified via ELISA. Hamsters who received intranasal boosts of CVXGA1 (group 2B), CVXGA13 (group 2C), or CVXGA14 (group 2E) had anti-S IgG titers greater than 5 log_10_, while hamsters who received a mRNA vaccine boost (group 2A) or no boost (group 2D) had lower anti-S IgG titers (Figure 7A). In neutralization assays with WA1, delta variant, and omicron variant, there was negligible difference in average neutralization titer between sera from hamsters who received a mRNA boost to hamsters who received no boost. However, boosting with mRNA increased detectable levels of neutralizing anti-WA1, delta and omicron from 4, 2 and 3 of 5 animals to 5, 3 and 5 of 5 respectively, indicating that a mRNA boost modestly increased neutralizing antibody responses (Figure 7B). In contrast, significantly higher levels of neutralizing antibodies against WA1, delta variant, and omicron variant were observed in hamsters boosted with CVXGA1, CVXGA13, or CVXGA14 (Figure 7B).

**Figure 7.**
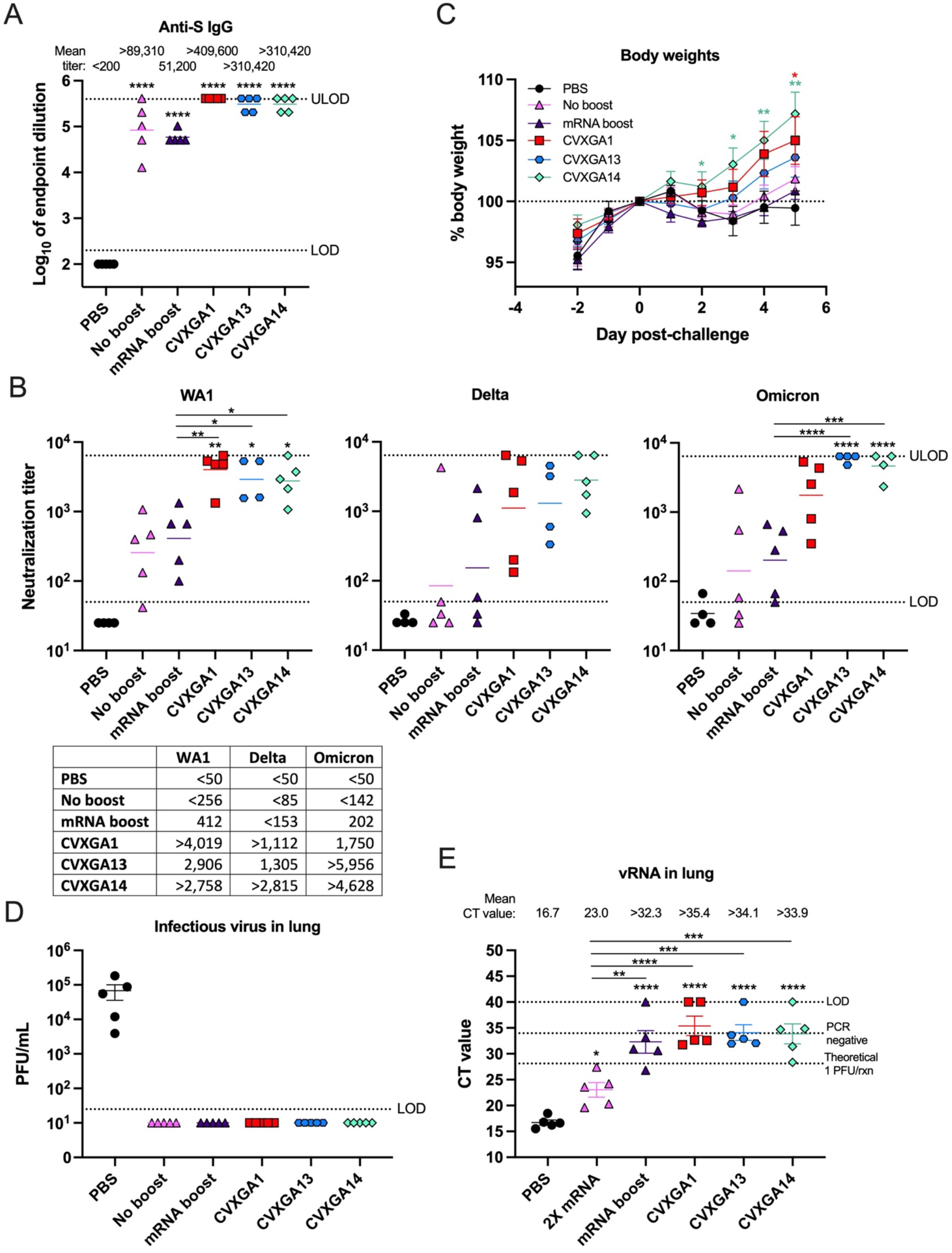
CVXGA1 boosts the humoral immune response and protection of mRNA-immunized hamsters against challenge with delta variant. (A) Anti-SARS-CoV-2 S IgG antibodies at 17 days post-boost were quantified via ELISA. Antibody titer was calculated as log_10_ of the highest serum dilution at which the OD_450_ was greater than two standard deviations above the mean OD_450_ of naïve serum. The limit of detection (LOD) is indicated by the dotted line. Bars represent geometric means. Statistical significance was calculated with one-way ANOVA (**** p < 0.0001). (B) Microneutralizing antibody titers against SARS-CoV-2 WA1, delta, or omicron were calculated as log_10_ of the highest serum dilution at which the virus infectivity was reduced by at least 50%. The limit of detection (LOD) is indicated by the dotted line. The geometric means are presented in the table and represented as bars on the graph. Statistical significance was calculated between each group and the PBS group or 2X mRNA group with one-way ANOVA (* p ≤ 0.05, ** p < 0.01, *** p < 0.001, **** p < 0.0001). (C) Hamster body weight changes over five days post-challenge were graphed as percent of day 0 weight. Statistical significance was calculated for each timepoint between each group and PBS-immunized hamsters with t tests (* p ≤ 0.05, ** p < 0.01). (E) Viral load in lung homogenates at 5 dpi was quantified via plaque assay in Vero TEMPRSS cells and graphed as PFU/mL lung homogenate. The limit of detection (LOD) is indicated by the dotted line. Error bars represent the standard error of the means. (F) SARS-CoV-2 delta vRNA load in lung homogenate was quantified via RT-qPCR. The cycle threshold (Ct) value for each sample is presented and error bars represent the standard error of the means. The known viral titer of delta variant was used to generate a standard curve and to calculate the Ct value equating to 1 PFU per reaction (rxn). The Ct value generated from RNA extracted from sterile water is denoted by a dotted line labelled PCR negative. The limit of detection (LOD) is indicated by a dotted line at Ct value = 40, the number of PCR cycles. Error bars represent the standard error of the means. Statistical significance was calculated with one-way ANOVA (** p < 0.01, **** p < 0.0001).

Twenty-five days following the boost, the hamsters were challenged with delta variant. Hamsters who received intranasal boosts of CVXGA14 or CVXGA1 had the best weight gain compared to hamsters received PBS, no boost, or an mRNA boost. Interestingly, hamsters who received CVXGA14 experienced higher weight gain than hamsters who received CVXGA13 (homologous delta S antigen) (Figure 7C). None of the boosted hamsters had detectable infectious virus in their lung homogenate at 5 dpc (Figure 7D; LOD, 25 PFU/mL). However, high levels of viral RNA were detected in animals with only two doses of mRNA. CVXGA-boosted animals had the least amount of viral RNA while mRNA-boosted animals had higher levels of viral RNA among all boosted groups (Figure 7E).

## Discussion

A PIV5-vectored SARS-CoV-2 vaccine expressing S (CVXGA1) has been shown to be efficacious against SARS-CoV-2 WA1 strain in mice and ferrets [17] and is currently in human clinical trials in the US. In this work, we demonstrated that one intranasal immunization of CVXGA1 protects hamsters against homologous WA1 and heterologous alpha and delta virus challenge (Figure 4, 5 and 6). Compared to 2X mRNA vaccine immunization, 2X CVXGA1 and 1X CVXGA1 generated lower anti-S binding antibody (Figure 6B), yet, anti-WA1 neutralizing antibody titers were similar among these immunization groups (Figure 6C). It is possible that because the S protein expressed by CVXGA1 is functional as indicated by their ability to promote syncytial (Figure 2), likely of native conformation (which may contain both pre- and post-fusion form of the S protein), CVXGA1 immunization may generate more cross-reactive and functional antibody responses than a mRNA vaccine. Seventy-two days post-boost immunization (at 108 dpi), anti-S antibody ELISA titers were similar among all the immunization regimens (Figure 6B). However, WA1-neutralizing antibody levels were highest in the 2X CVXGA1 group and lowest in the 2X mRNA group, indicating that CVXGA1 immunization maintains neutralizing antibody levels better than 2X mRNA immunization. The rapid reduction of neutralizing antibody titers in hamster following 2X mRNA vaccine immunization (Figure 3C) is consistent with reports of rapid reduction of neutralizing antibody titers after two doses of mRNA vaccine immunization [30]. The long-lasting neutralizing antibody levels from CVXGA1 immunization (Figure 6B and Figure 6C) may be attributed to the intranasally-expressed S antigen delivered by the live replicating PIV5 vector. While a single dose intranasal immunization with CVXGA1 protects hamsters against WA1 or VOCs alpha and delta, 2X CVXGA1 generates longer-lasting immunity (Figure 6C), better body weight gain after challenge (Figure 6D), and lower viral load as judged by viral RNA levels after challenge (Figure 6F), indicating that boosting CVXGA1-immunized animal with CVXGA1 affords additional protection.

Most of the US population has received at least one COVID-19 immunization (https://covid.cdc.gov/covid-data-tracker). Fully vaccinated individuals have omicron-neutralizing antibody titers 22-fold lower than WA1-neutralizing antibody titers [31] and individuals having received three doses of COVID-19 mRNA vaccines BNT162b2 or mRNA-1273 have omicron-neutralizing antibody titers of 10.7- and 7.2-fold lower, respectively [32]. It has been reported that heterologous prime-boost generates more robust immune responses than homologous prime-boost [33, 34]. Therefore, we determined the immunogenicity and efficacy of CVXGA1 as a booster in hamsters immunized with 2X mRNA vaccine (Figure 7A and 7B). As expected, boosting 2X mRNA-immunized hamsters with a third dose of mRNA vaccine did not significantly increase anti-S binding antibody titers (Figure 7A) and only moderately increased neutralizing antibody titers (Figure 7B). In contrast, boosting 2X mRNA-immunized hamsters with an intranasal dose of CVXGA1 resulted in an approximate 4.6-fold increase of anti-S antibody levels (Figure 7A) and a significant increase of neutralizing antibody titers (>15.7-, >13.1-, and >12.3-fold for WA1, delta, and omicron, respectively) (Figure 7B). Comparing mRNA vaccine-boosted hamsters (three doses of mRNA vaccine), hamsters boosted with CVXGA1 (2X mRNA plus one dose of CVXGA1) had more body weight gain (Figure 7C) and lower viral load as judged by RT-qPCR of the lungs of challenged animals (Figure 7E), indicating that boosting with CVXGA1 resulted in better outcome for hamsters than boosting with a mRNA vaccine. These results are consistent with human studies for heterologous vaccine prime-boost [33, 34].

While we confirmed the benefits of a heterologous prime-boost with different vaccine platforms, we also observed the advantages of a heterologous antigen prime-boost. Intriguingly, we found that boosting 2X mRNA-immunized hamsters with a viral vector expressing delta (CVXGA13) or omicron S (CVXGA14) substantially increased neutralizing antibody levels against all three strains WA1, delta variant, and omicron variant. Furthermore, boosting 2X mRNA vaccinated animals with CVXGA13 or CVXGA14 increased WA1-neutralizing antibody titers significantly better than a homologous mRNA vaccine boost (Figure 7B). The differences in antigen presentation and vaccine delivery route (intranasal for CVXGA vaccines vs. intramuscular injection for mRNA vaccines) may have contributed to the more robust antibody responses induced by a CVXGA vaccine booster.

While we did not detect a clear advantage of using PIV5-vectored variant S vaccines over the ancestral S-based vaccine in alpha and delta VOC challenge in hamsters, the PIV5 vector has the capability for “plug and play” to quickly replace the target antigen. Due to the lack of pathogenicity in the hamster model, omicron was not used in our studies as a challenge virus [22]. VOC delta, to whom WA1-based mRNA vaccine has lower cross-reactive neutralizing antibodies against and causes the most body weight loss in hamsters, was chosen as the main challenge virus in our studies. It will be interesting to test CVXGA14, or additional PIV5-vectored vaccine constructs, against yet-to-emerge SARS-CoV-2 variants in the future. Finally, SARS-CoV-2 N-specific immune responses may be protective. CVXGA2, expressing both S and N antigens of SARS-CoV-2 WA1, did have the most body weight gain and lowest viral load after heterologous virus challenge (Figure 4B and 4F). Thus, it appears that the N protein might offer additional protection, and its mechanism of protection will be further evaluated.

Due to the experimental limitation, we did not measure mucosal immunity or cellular immunity in these studies. Intranasal immunization with a live viral vector is expected to result in mucosal and cellular immunity in addition to humoral immune responses [12]. In this study, we did not detect a direct correlation between neutralizing antibody titers against delta and delta viral load, suggesting that other immune responses, such as cellular immune responses and/or mucosal immunity, might have played critical roles in protection.

In summary, CVXGA1, an intranasal vaccine currently being evaluated in human clinical trial in the US [23], protects against challenge with the homologous SARS-CoV-2 virus strain and heterologous VOCs as a single-dose in naïve and in mRNA-immunized hamsters. Our data suggests that CVXGA1, and other CVXGA vaccines, can serve as an effective heterologous booster to offer longer-lasting immunity to those who have received COVID-19 mRNA vaccines.

## Acknowledgements

We thank members of Drs. Biao He’s, Mark Tompkins’s, Jeff Hogan’s, and Eric Lafontaine’s labs for their technical support and helpful discussions. The work by Dr. Michael Gale, Jr. was supported by NIH grant AI151698 (MG). We thank the UGA Animal Resources staff.

## Supplemental figure

**Supplemental figure 1.**
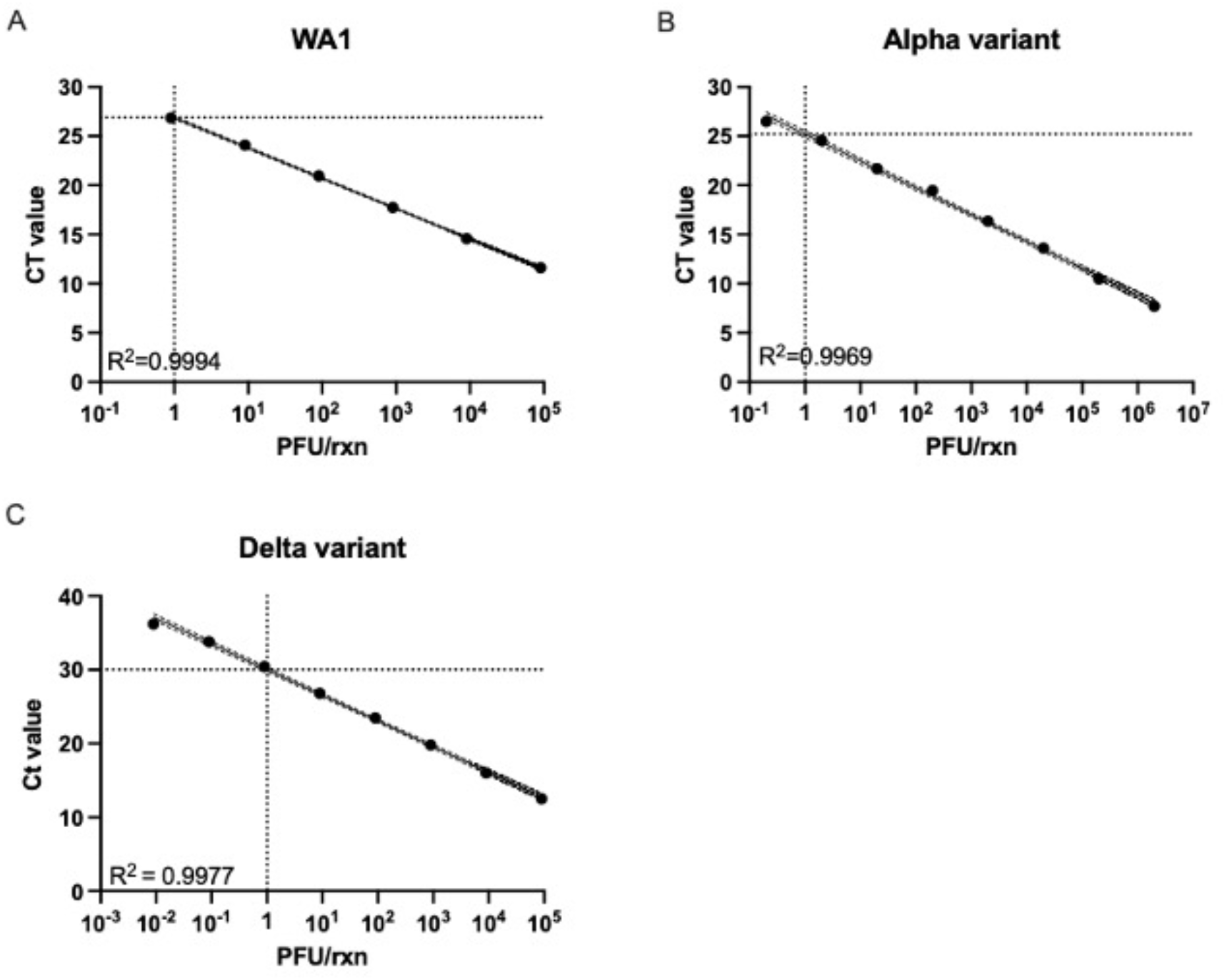
RT-qPCR standard curves. RNA was extracted from SARS-CoV-2 WA1 (A), alpha variant (B), and delta variant (C) viral stocks of known titer. The RNA was serially diluted and vRNA was quantified via RT-qPCR. To generate a standard curve, the Ct value was plotted on the y-axis and the PFU per reaction (rxn) was plotted on the x-axis. Dotted lines indicate the Ct value which corresponds to 1 PFU/rxn.

